# Identification of bioactive constituents for colitis from traditional Chinese medicine prescription via deep neural network

**DOI:** 10.1101/2023.05.31.542690

**Authors:** Zhixiang Ren, Yiming Ren, Pengfei Liu, Qi Shu, Huijuan Ma, Huan Xu

**Author notes:** Correspondence (H. Ma); (H. Xu).

## Abstract

Colitis is a commonly encountered inflammatory disease in colon tissue, which can be triggered by various causes. Although traditional Chinese medicine (TCM) has been utilized for the treatment of colitis, it is still a great challenge to identify the major bioactive constituents and their modes of action among thousands of ingredients in TCM prescriptions. Inspired by the success of artificial intelligence and deep learning methods, we proposed a deep neural network (DNN) for TCM prescription recommendation. We constructed a graph-based DNN with 9,845 nodes and 161,950 edges, which integrated microscopic information including bioactive molecules, protein targets, and extracted features of prescriptions through feature embedding. A novel and efficient data augmentation strategy was implemented to expand the sample size based on 378 collected TCM prescriptions. Network pharmacology study revealed that the 10 most frequent ingredients in generated prescriptions were associated with multiple inflammatory signaling pathways. To verify the bioactive constituents in the generated prescriptions, 5 selected constituents were administrated to BALB/c mice with colitis. Suppressive effects of disease progression and pro-inflammatory factors comparable to sulfasalazin were observed with these compounds, revealing the effectiveness of our artificial intelligence strategy on idetification of bioactive constituents from TCM prescriptions.

## I. INTRODUCTION

**I**NFLAMMATORY bowel disease (IBD) encompasses a wide range of intestinal inflammatory conditions, such as the chronic relapsing Crohn’ s disease and ulcerative colitis, is becoming increasingly common in China [1]. The pathogenic mechanisms of IBD are quite complicated, which involves the interactions between genetic, environmental and microbial factors [2]. Besides the IBD colitis, other forms of inflammatory conditions, including infectious colitis, microscopic colitis, ischemic colitis, segmental colitis, drug-induced colitis and diversion colitis, are also prevalent and sometimes agnogenic [3]. Current therapeutic agents for colitis include aminosalicylates, corticosteroids, immunosuppressants, cyclosporine and monoclonal antibodies [4]. However, most of these compounds trigger various adverse reactions and cannot produce long-lasting treatment effects [5]. Therefore, it is of great significance to develop more efficient drugs for colitis treatment. Traditional Chinese medicine (TCM) is based on distinctive Chinese cultural theories and thousand years’ clinical practices, which has been utilized to treat colitis and produced many successful herb formulas [6]. These formulas can be found in different TCM books, such as Shennong’ s Classic of Materia Medica, Reatise on Cold Pathogenic and Miscellaneous Diseases, Prescriptions for Universal Relief and so on. Although the treatment effects of the formulas have been verified by clinical practices, their major bioactive constituents and molecular mechanisms have not yet been identified.

The diagnoses of TCM are mainly based on systematic symptoms or signs obtained by observation, listening, asking and cutting. Prescriptions are usually sets of herbs, which may be precise and varied for individuals carrying same symptoms [7]. Over the years, many successful nature compounds had been derived from TCM formulas for the treatment of diseases, such as artemisinin, tetrahydropalmatine and tetramethyl-pyrazine [8]. With the establishments of multiple herb-ingredient databases, such as the Traditional Chinese Medicine Systems Pharmacology Database and Analysis Platform (TCMSP), now it takes little effort to figure out the ingredients and molecular targets of a single herb [9]. However, based on the number and complexity of components in prescriptions, it is a great challenge to identify the active compounds among thousands of constituents. In this case, novel in silico tools, such as the machine learning-based methods and network pharmacology, has been utilized to uncover the bioactive constituents and molecular targets behind the herb formulas. For examples, Dong and colleagues built a subnetwork of herb-symptom for recommendation of TCM prescriptions [10], while Zhou et al. integrated disease phenotypes and constituents into a deep neural network (DNN) for formula recommendations [11]. On the other hand, network pharmacology through data mining has also been verified to be useful in finding bioactive molecules from TCM formulas for certain diseases [12,13]. Both of them can be utilized for accurate and efficient identification of the bioactive constituents for certain diseases.

In this study, we proposed a graph-based DNN that recommends TCM prescriptions by integrating information from TCM formulas and bioactive molecules. Initially, a graph network contains herb-ingredient-target relationships was constructed using open-source datasets, such as TCMSP and PubChem. We modeled the correlation between herbs and bioactive molecules by employing graph embedding to extract the latent features of TCM formulas. Furthermore, a DNN was introduced to evaluate the effectiveness of TCM prescriptions, which assigned a score ranging from 0 to 1 to candidate prescriptions. To mitigate overfitting, we designed a data augmentation strategy that expanded the training dataset by augmenting real TCM formulas acquired from ancient books, based on prior knowledge. Finally, we inferred all possible combinations of 2, 3, and 4 herbs using this model and selected the top-10 effective prescriptions for subsequent biological experimental validation. The intersected bioactive constituents in the recommended herb formula with highest score and the most frequent set of ingredients identified in network pharmacology study were tested in dextran sulfate sodium salt (DSS)-induced murine colitis models to verify their treatment effects on colitis. The results not only confirmed the performance and quality of TCM formula recommendation through neural network for colitis, but also provided valuable information for the future development of therapeutic natural compounds from TCM formulas.

## II. MATERIALS AND METHODS

### A. Data source and processing

### 1) Augmentation of real-world prescriptions

Real-world prescriptions for colitis were firstly excerpted from 204 TCM books or databases. A total of 378 distinct prescriptions were collected, which was insufficient to train a high-performance neural network. Therefore, a data augmentation strategy was introduced for dataset generation. The highest evaluation score was awarded to the real-world prescriptions [11], while the generated prescriptions were assigned with a lower score (<1) based on the similarity to the real-world prescriptions, which was measured by Jaccard coefficient combined with prior information:

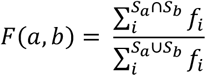

As shown in the formula, *s*_a_ and *s*_b_ respectively represented the herb sets of two distinct prescriptions, whereas f_i_ was a set containing the occurrence frequency of each herb in the real prescriptions. The similarity F (a, b) was defined as the ratio of the sum of the frequencies of the intersection of a and b to the sum of the frequencies of the union of a and b. In the data augmentation phase, we randomly sampled herbs from the real-world prescriptions and calculate their similarity score. As a result, a total of 85,583 formulas were generated for neural network training, and the general distribution of dataset was shown in Figure 1.

**Figure. 1.**
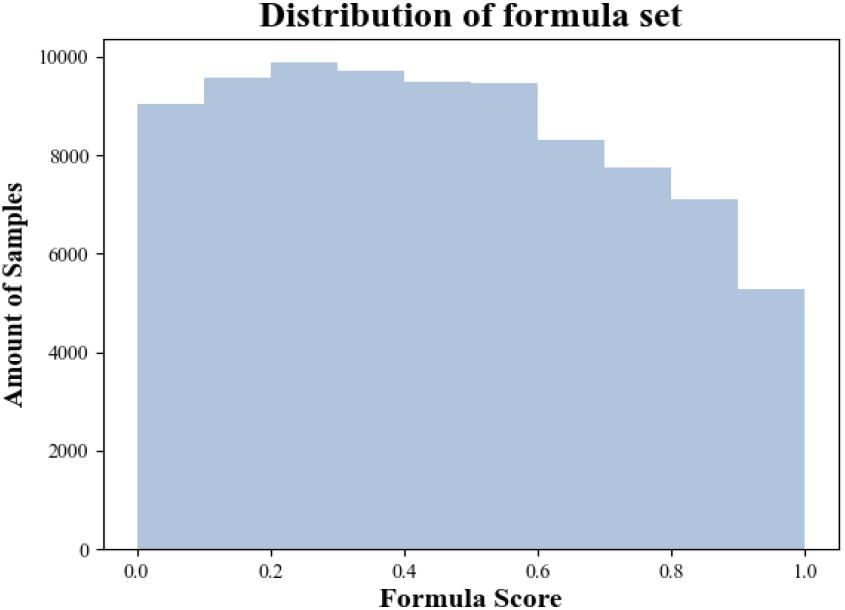
The score distribution of the generated TCM formula dataset with the amount of 85,583 samples augmented from 378 distinct real-world prescriptions. With 0.1 as the unit interval, the distribution of data in each interval was uniform.

### 2) Graph construction

The interactions of herbs, ingredients, and targets were modeled into a graph network. The information in TCMSP (https://old.tcmsp-e.com/tcmsp.php) was utilized to determine the relationships between herb and primary constituents or constituents and protein targets. Additionally, the similarity between constituents were captured from PubChem, and only constituent pairs with a similarity greater than 90 percent were concerned. Eventually, the large-scale graph network was constructed with 9845 nodes, containing 218 herbs, 7987 constituents and 1640 protein targets. The amounts of edges were 161950, which included linkages of 18912 herb-constituent, 101651 constituent-constituent and 41387 constituent-target.

### B) Neural Network Architecture

The architecture of neural network was summarized in Figure 2, including the graph embedding layer and herb recommendation layer. We implemented the graph embedding layer according to the PyTorch Geometric, which is a library for deep learning on graph data [14]. Based on the nature of herb-constituent-target network, Node2Vec was utilized to capture the latent features of each herb. Node2Vec is a classic algorithm for graph embedding that can be used in large-scale networks [15]. The objective of the algorithm was to maximize the similarity between two nodes with a similar context. Therefore, each node was first initiated to a matrix with a predetermined dimension and the matrix of each node aggregated its neighborhood information through multiple messages passing. The objective function of Node2Vec was:

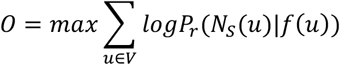

The set of the neighborhood *N*_*s*_(u) of node u was generated with a certain sampling strategy. f(u) represented the function that maps the node u to the embedding matrix. The objective function was to maximize the probability P*s*_*r*_ of sampling the actual neighboring nodes in order to determine the precise similarity between nodes. The relationships of constituents and targets corresponding to each herb were considered as context. Therefore, the algorithm was able to mechanistically assess the herb with the highest similarity and then assist the TCM formula recommendation. On the other hand, the herb recommendation layer of the proposed model involved a multi-layer perception (MLP) network, a widely used neural network architecture for classification and regression tasks. The input to this layer was the feature matrix of TCM formula, which was combined with the feature matrices of herbs extracted from the graph embedding layer.

**Figure. 2.**
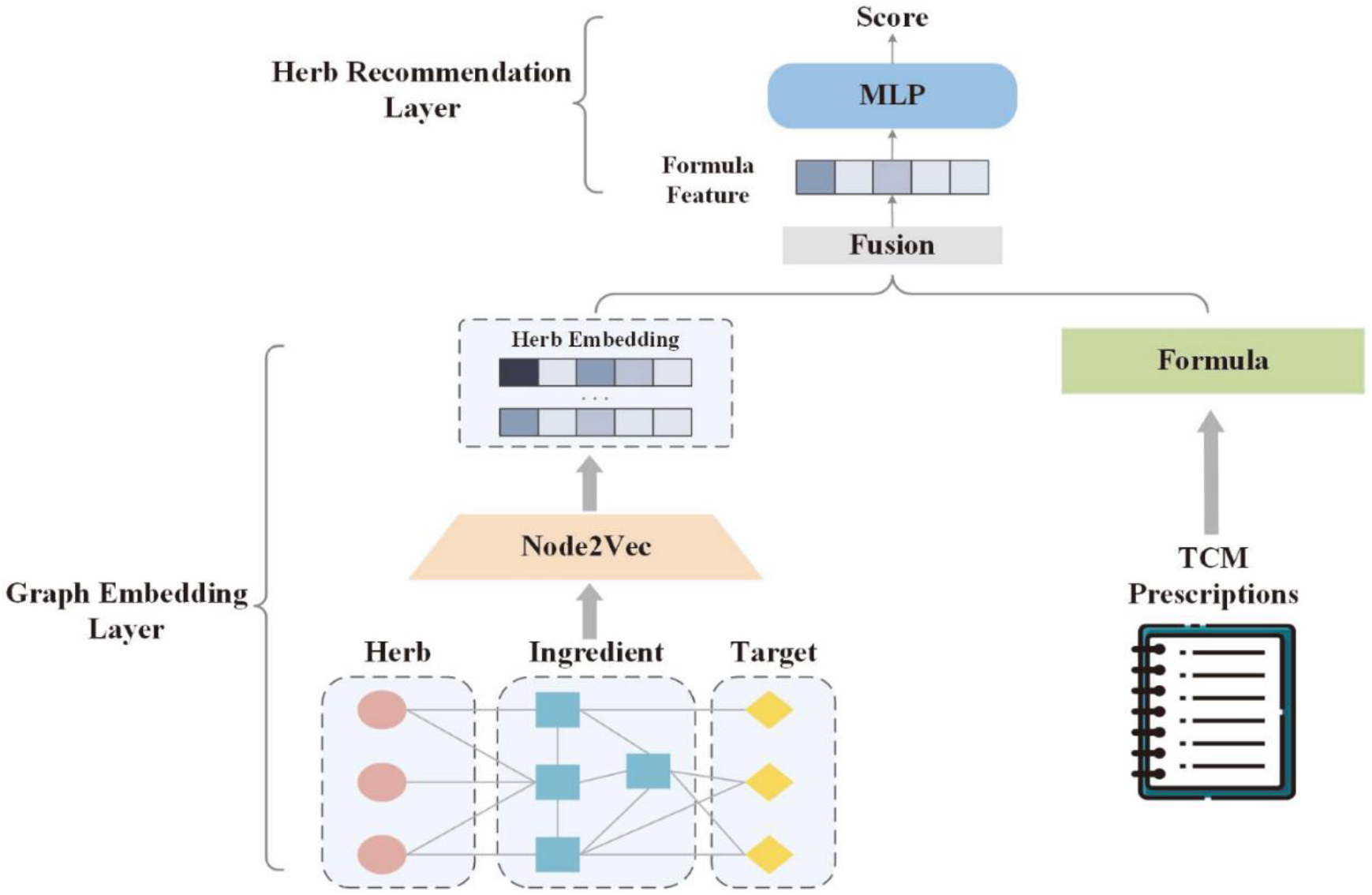
The architecture of neural network model. The graph embedding layer was used to extract the latent feature of herb from the network. The herb recommendation layer evaluated the quality of the prescriptions by fitting the feature of each formula.

The formula feature matrices were measured by averaging the respective combination herb matrix. The MLP was trained to regress the formula feature matrices and generate a score for each formula. The MSE loss was used as the loss function to perform global optimization, where 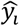 and *ys*_*i*_ represented the predicted and real scores, respectively:

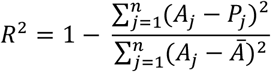

By minimizing the loss, the herb recommendation layer was able to accurately predict the quality of the TCM formulas.

### C. Experimental settings of neural network

In the experimenting phase, we trained the model within the two parts separately. The training process of Graph Embedding was presented in Figure 3A, in which the loss began from 6.67 and converged to 0.72 by the 250th epoch. In the herb recommendation module, we divided the dataset into the training set, the test set and the validation set in a ratio of 8:1:1. The validation set was applied to prevent the model from overfitting. R-square score (*Rs*^2*s*^) was utilized as the evaluation metric, which was defined as follows:

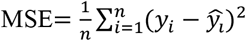

*R*^2^ was the coefficient of determination, which indicated the proportion of variation in the dependent variable that could be predicted. *A*_*j*_ was the ground truth, *P*_*j*_ was the predicted value and Ā represented the mean of ground truth. The R-square score of the train set was depicted by the blue curve, which hits 0.957 on the 124th epoch. The orange curve represented the R-square score on the test set, which finally reached 0.954, validating the high performance of the model (Figure 3B).

**Figure. 3.**
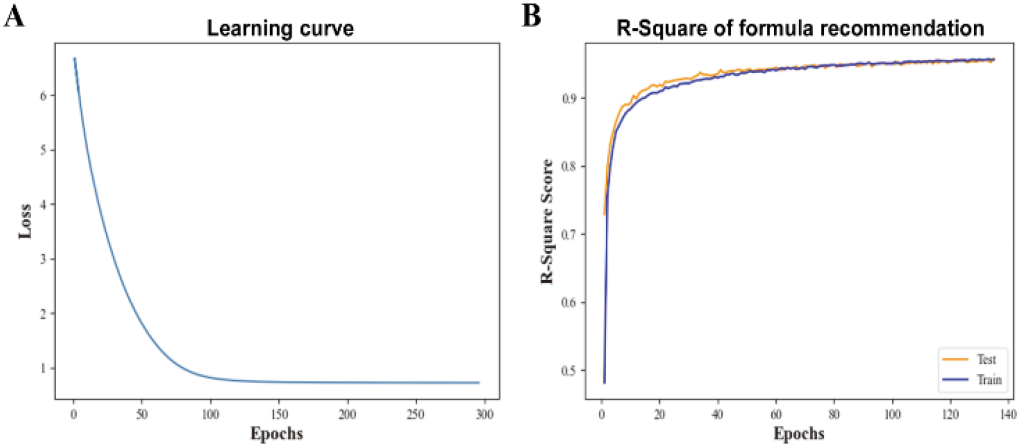
The learning curve of the graph embedding layer and the R-square score of the herb recommendation layer. A, the learning curve indicated a relatively steady convergence of the training process. B, the R-square score, ranged from 0 to 1, was utilized to measure the regression accuracy.

### D) Network pharmacology study based on the bioactive constituents

All bioactive constituents in the 378 prescriptions and their corresponding targets were retrieved from TCMSP database (see supplemental materials 1). The frequency of each constituent in all prescriptions were sorted from high to low, and the molecular targets of the 10 constituents with highest counts were converted to gene names and compared with the colitis-associated genes acquired from DisGeNET (https://www.disgenet.org/, CUI: C0009319). The gene ontology (GO) analysis of the gene intersections was performed on the Database for Annotation, Visualization and Integrated Discovery (DAVID, https://david.ncifcrf.gov/), and the identified pathways were graphed. The constituent-target network was constructed using Cytoscape 3.7.2 and the protein-protein interaction (PPI) network was generated on STRING with the intersected genes (https://cn.string-db.org/).

### E) Murine model of colitis and treatments

Male BALB/c mice were purchased at 7 weeks of age from JSJ-Lab (Shanghai, China). After 1 week of acclimation in our animal facility, mice were randomly divided into 8 groups (n = 6, a total of 48 mice). Three mice in the same group were caged together on corn cob bedding, with standard chow (NovoBiotec, Beijing, China) and boiled tap water ad libitum. DSS (CAS 9011-18-1, mol wt ∼40,000, Sigma-Aldrich, St. Louis, MO, USA) was added to the drinking water at 3% to induce colitis in 8 d (Figure S1 in supplemental materials 2). Disease progressions were monitored by body weights and stool consistencies during the DSS treatment period. Daily gavage administrations of the identified bioactive constituents, including β-sitosterol (CAS 83-46-5, Sigma-Aldrich), quercetin (CAS 117-39-5, Sigma-Aldrich), (R)-linalool (CAS 126-91-0, Sigma-Aldrich), myristic acid (CAS 544-63-8, TCI), oleanolic acid (CAS 508-02-1, Sigma-Aldrich) and sulfasalazine (SASP, CAS 599-79-1, TCI, used as the positive control) were conducted from day 9 to 15 to alleviate the inflammatory responses of colitis. Serum and colon samples were collected after sacrifice by cervical dislocation at the end of the experimental period.

### F) Inflammatory cytokine, MPO and iNOS assays

The cytokine expressions were assayed by ELISA kits of mouse IL-1β, TNF-α, IL-6, IL-18 and MCP-1 (Abcam, Cambridge, UK). Serum samples were diluted 1:10 and loaded to the strip wells precoated with primary antibodies. After adding the cocktail with HRP-conjugated secondary detector antibody, the microplates were sealed and incubated on a plate shaker set at 400 rpm for 2 h at RT. The TMB chromogen solution was then added to the wells and the absorbance was recorded at 450 nm on a microplate reader for calculation of cytokine concentrations based on the standard curves. For the MPO assay (Abcam), homogenized colon tissue samples were quantified by BCA assay and mixed with MPO substrate in the assay buffer in a 96-well microplate. After 1 h of incubation at RT in dark, the stop mix and TNB reagent were added to the wells and 412 nm absorbance were recorded on a microplate reader. For the examination of iNOS activity in colon macrophages, tissue samples were carefully trimmed into tiny pieces with a sterile iris scissor and digested by 100 U/mL collagenase I and 2.4 U/mL dispase II (Yeasen, Shanghai, China) at 37 °C on a shaker set at 100 rpm for 2 h. The digested samples were filtered with sterile 40 μm cell strainers, centrifuged for 5 min at 400 ×g, and resuspended in PBS buffer with 1 μg/mL biotin-labeled F4/80 antibody (Abcam). After 30 min incubation, macrophages were then incubated with SAV magBeads for 30 min at RT and retrieved with a magnetic separation rack. The reaction buffer of the iNOS activity kit (Beyotime, Shanghai, China) containing L-Arginine, NADPH and DAF-FM DA (NO probe) was then added to the cells in a Corning 96-well black microplate with clear flat bottom, and the fluorescence was acquired on a microplate reader (Ex/Em 495/515 nm) for the calculation of iNOS activity in samples according to the user manual provided by the manufacturer.

### G) Statistics

Data generated in the cell and animal experiments was analyzed with Excel 2016 and GraphPad Prism v8.0.1. For the murine model of colitis, 6 mice were used in each group. For the cell viability study using human colonic epithelial cells (HCoEpiC, ScienCell, third passage), 3 independent experiments were performed and analyzed for each individual concentration of the treatments. One-way analysis of variance (ANOVA) and Turkey’ s post-hoc tests were applied to determine differences between groups. P values under 0.05 were considered statistically significant.

## III. RESULTS

### A. Generation of high quality TCM formulas for colitis via neural network

After the construction of TCM pharmacological neural network based on the 378 TCM colitis formulas following augmentation, the trained model was utilized to recommend high-quality herb combinations for generation of new prescriptions. The network parameters learned by the Graph embedding layer and the Herb recommendation layer were respectively retained for further operations. Prior to the inference process, we loaded the embedding matrices of each herb and calculated the feature matrices of all candidate formulas, which served as the inputs for recommendation process. The weights of herb recommendation network were loaded to infer the scores of all candidate formulas. Due to the stochastic nature of the DNN, the prescription scores obtained in each run slightly changed by no more than 0.0006, thus we performed multiple runs to infer prescription scores, and averaged the outputs to achieve relatively stable results. We generated TCM formulas composed of 2, 3, or 4 herbs using all possible combinations of 218 herbs, totaling 93,263,779 groups, and identified the top 5 highest-scored formulas in each combination (Table 1). Since the formulas of 4 herbs achieved higher scores, the ingredients of the highest-scored 4 herb formula were utilized in the following studies.

**Table I.**
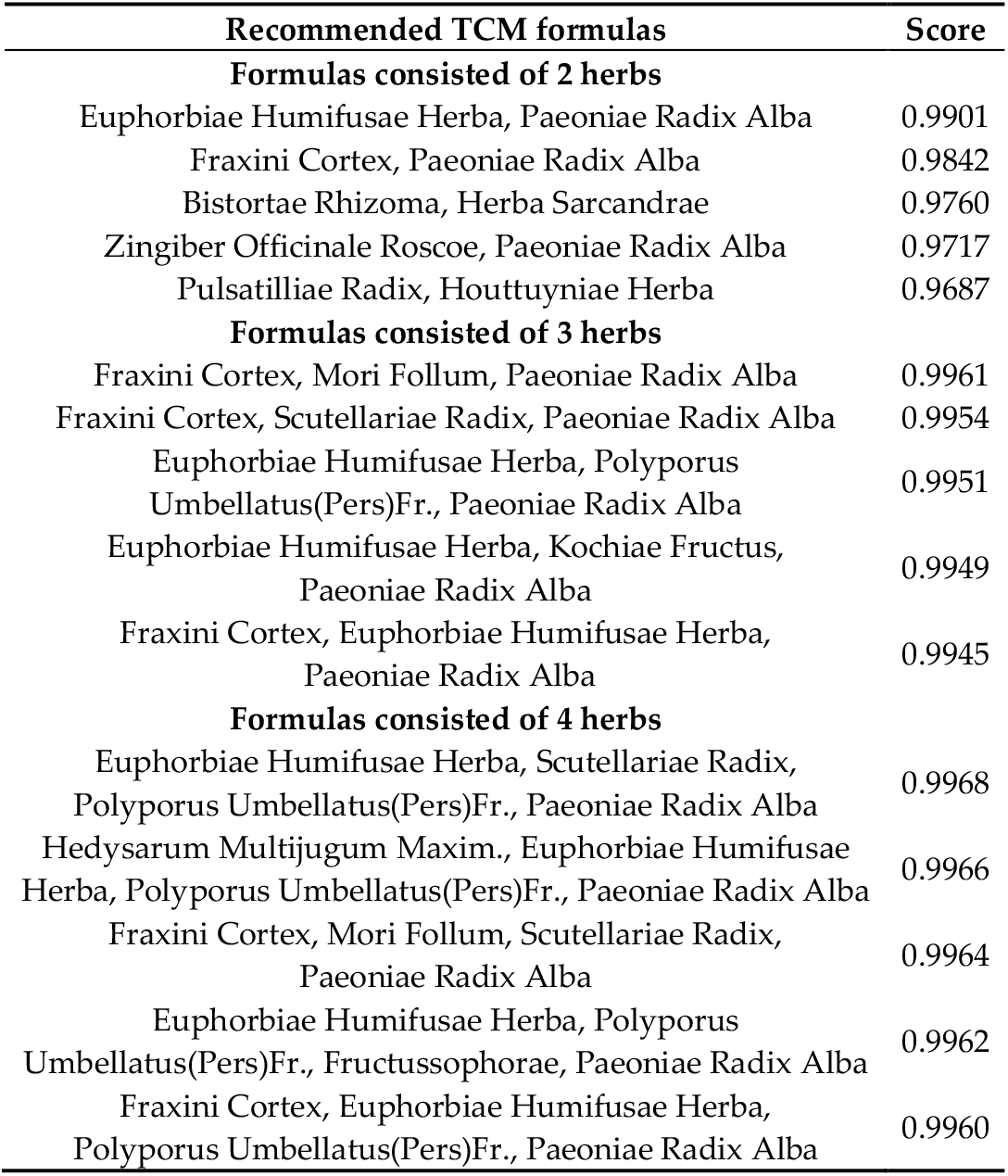
TOP Recommended TCM formulas

### B. Network pharmacology analysis of major bioactive constituents for colitis from the TCM formulas

After the ranking of all the 7987 ingredients in the herbs of 378 TCM colitis formulas based on their frequencies, the top 10 bioactive constituents were identified for network pharmacology analysis (Figure 4A). Among the genes encoding the 210 protein targets of the 10 constituents, 102 (48.6%) were intersected with colitis-associated genes (Figure 4B). KEGG gene enrichment analysis showed that the intersected genes were enriched into signaling pathways deeply involved in inflammation, such as AGE-RAGE, HIF-1, TCR and inflammatory cytokine pathways (Figure 4C), which was consistent with the biological process (BP), cellular component (CC) and molecular function (MF results) analyzed on PANTHER (Figure S1 in supplemental materials 2). Through the construction of constituent-target network, it became obvious that genes of transcriptional factors, tyrosine kinases, cytokines and metabolic enzymes involved in colitis were targeted by the molecules (Figure 5A). The PPI network further demonstrated that many inflammatory factors, such as the inflammation biomarkers MPO and CRP, were potential targets of the 10 constituents (Figure 5B). These results from the network pharmacology study indicated that the 10 bioactive constituents identified by their frequencies could target the inflammatory responses in colitis.

**Figure. 4.**
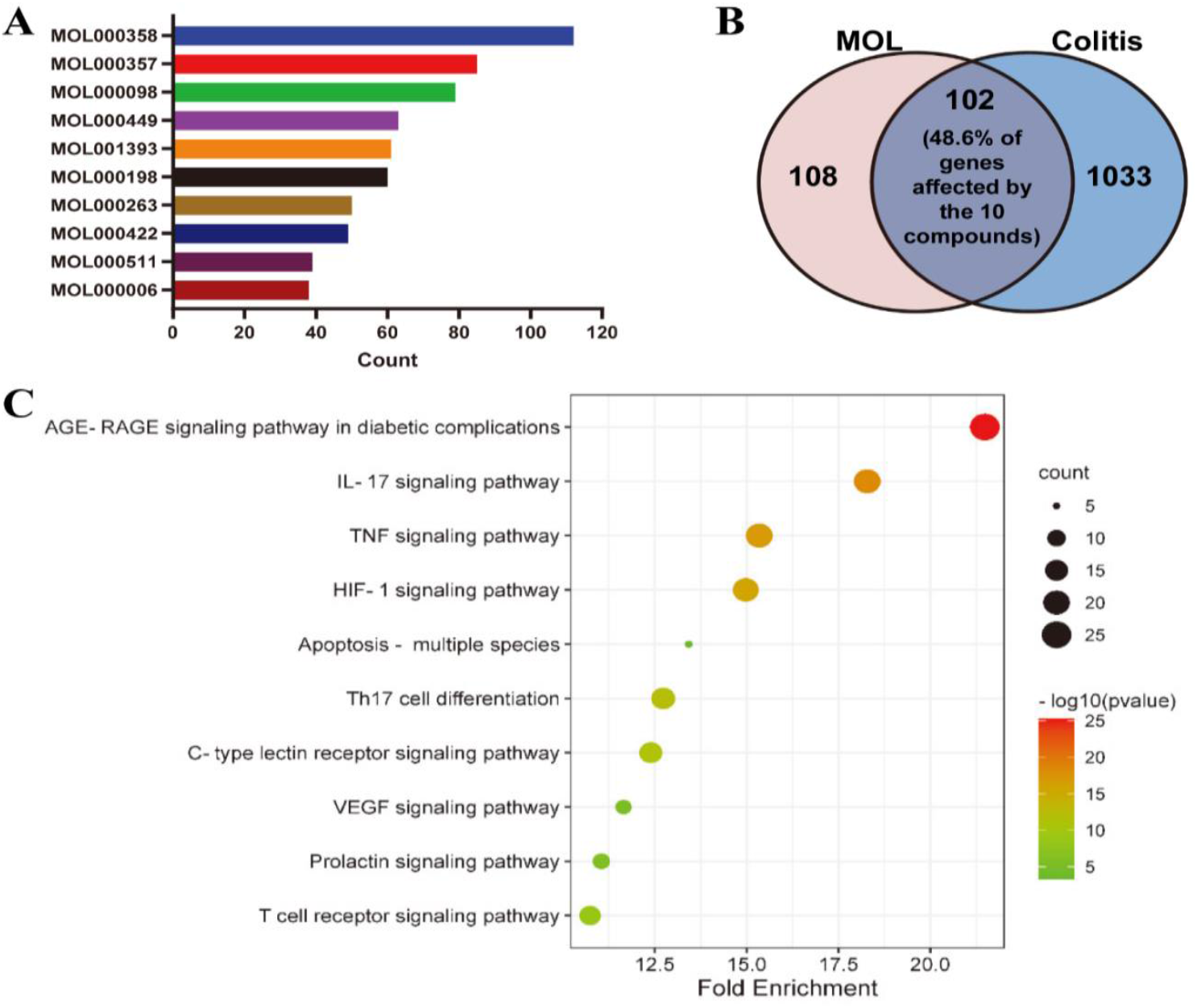
Targets and BPs from gene enrichment analysis. The constituents in each herb were acquired from TCMSP and their frequencies were sorted from high to low. The protein targets of the 10 constituents with highest counts were converted to gene names and compared with the colitis-associated genes acquired from DisGeNET. GO analysis was performed on DAVID based on the intersected genes. A, top-ranked constituents and their frequencies in the prescriptions. B, a venn graph showing the number of intersected genes. C, BPs enriched by the intersected genes ranked by fold enrichment.

**Figure. 5.**
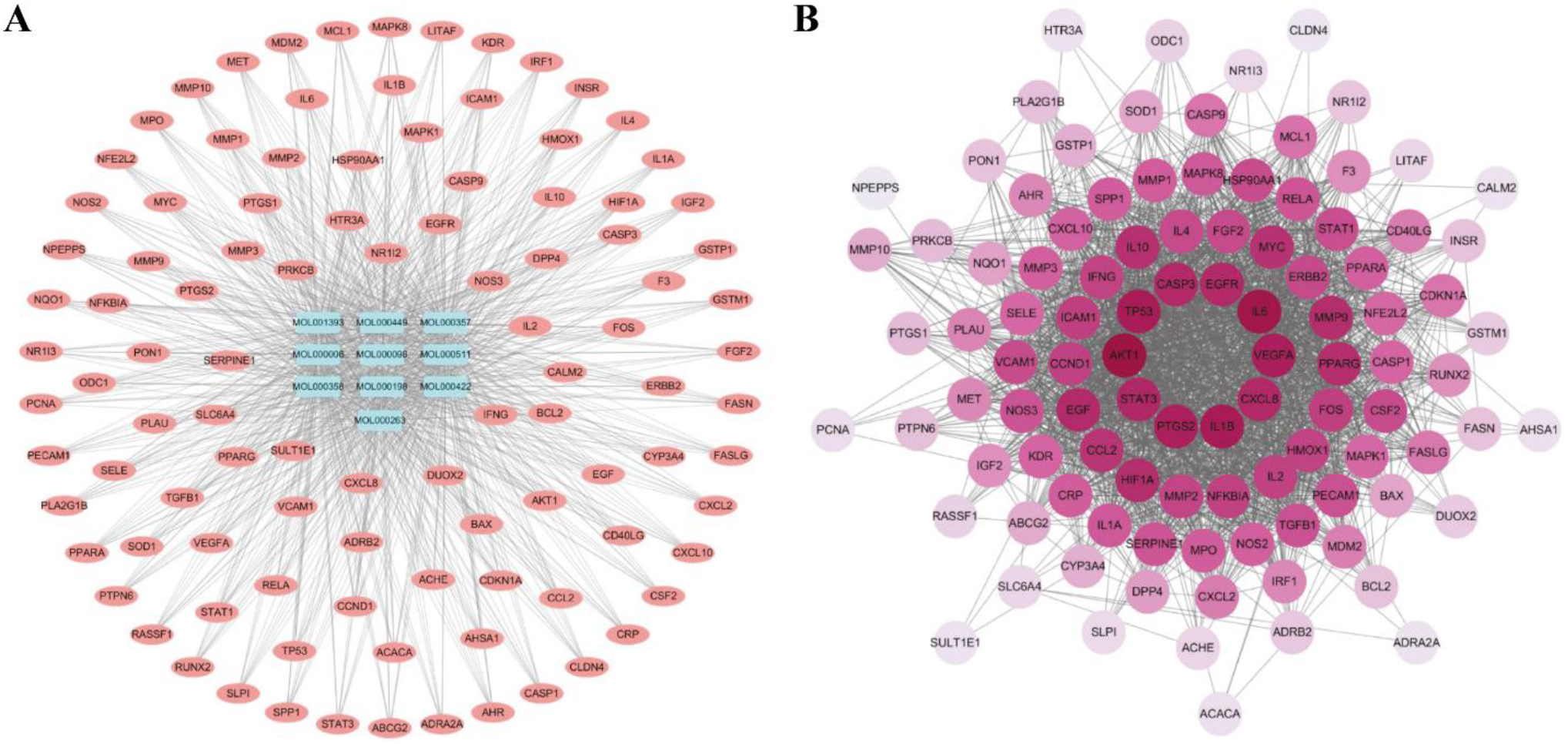
MOL-target and PPI networks. After identification of the 10 constituents with highest counts, the constituent-target network was constructed using Cytoscape 3.7.2. The protein-protein interaction (PPI) network was generated on STRING using the intersected genes. A, the constituent-target network. B, the PPI network.

### C. Treatment effects of the bioactive constituents in DSS-induced murine model of colitis

To verify the therapeutic effects of the compounds identified by TCM formula generation and network pharmacology, colitis was induced in mice via 3% DSS supplement in drinking water. The 5 constituents in the intersection between the formula’ s ingredients and the 10 most frequent compound in all real-world colitis formulas, including β-sitosterol, quercetin, (R)-linalool, myristic acid and oleanolic acid, were tested for colitis treatment (Figure 6A and B). After the examination of cytotoxicity via CCK-8 assays in HCoEpiC normal human colon epithelial cells (Figure 6C), β-sitosterol (100 mg/kg), quercetin (50 mg/kg), (R)-linalool (10 mg/kg), myristic acid (100 mg/kg), oleanolic acid (10 mg/kg) and SASP (100 mg/kg, positive control) were administrated daily through gavage to alleviate the inflammatory responses induced by DSS in colon. The DSS-induced decreases in colon length were significantly alleviated by all the other constituents except (R)-linalool (Figure 7A). The severity of colitis was also measured by disease activity index (DAI), which was the combined score of body weight and stool consistency (Table S1 in supplemental materials 2) (Zhang et al., 2020). All the 5 constituents reduced DAI of colitis, demonstrating their effectiveness in treatment of colitis (Figure 7B). Attenuation of inflammatory conditions were also observed with the treatments in the histopathological images [16]. In addition, serum levels of inflammatory cytokines, including IL-1β, TNF-α, IL-6, IL-18 and MCP-1, were reduced by the treatments in the mice with colitis (Figure 8A). The DSS-induced elevations of MPO activity in colon tissues and iNOS activity in colon macrophages were also reversed by the compound administrations (Figure 8B and C). Collectively, these results not only confirmed the therapeutic activities of the identified constituents, but also indicated that these natural products could be further developed as therapeutic agents for colitis.

**Figure. 6.**
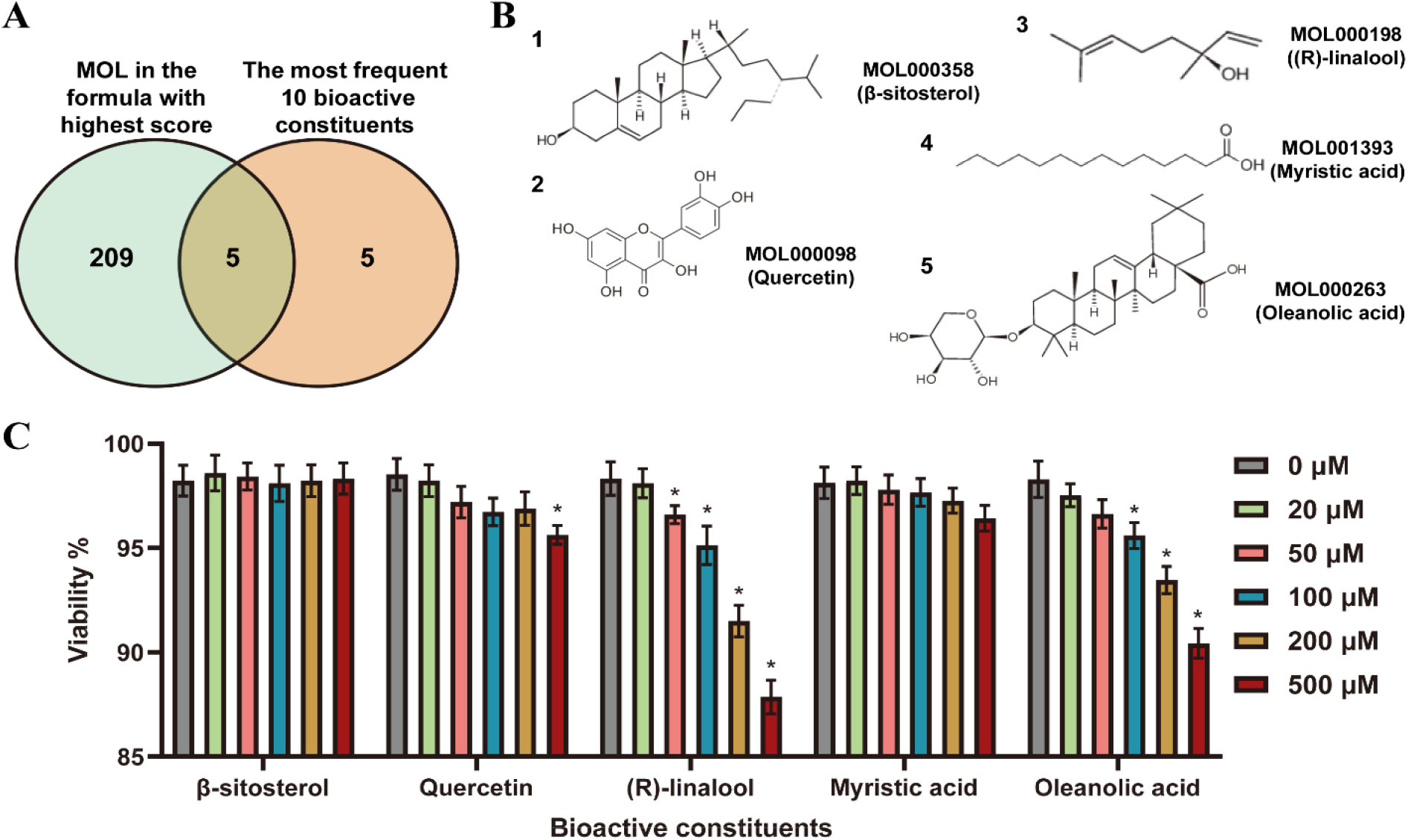
Bioactive constituents identified by neural network and network pharmacology. The 5 constituents in the intersection between the ingredients in highest-scored formula and the 10 most frequent compound in all real-world colitis formulas, including β-sitosterol, quercetin, (R)-linalool, myristic acid and oleanolic acid, were selected for the treatment of mice with DSS-induced colitis. The cytotoxicity of these compounds was analyzed via CCK-8 assay in HCoEpiC cells. A, a venn graph showing the number of intersected constituents. B, chemical structures of the 5 constituents. C, cell viability of HCoEpiC cells treated with the 5 constituents for 24 h in vitro. Results are mean ± SD. *Significantly different from the 0 μM group (p < 0.05).

**Figure. 7.**
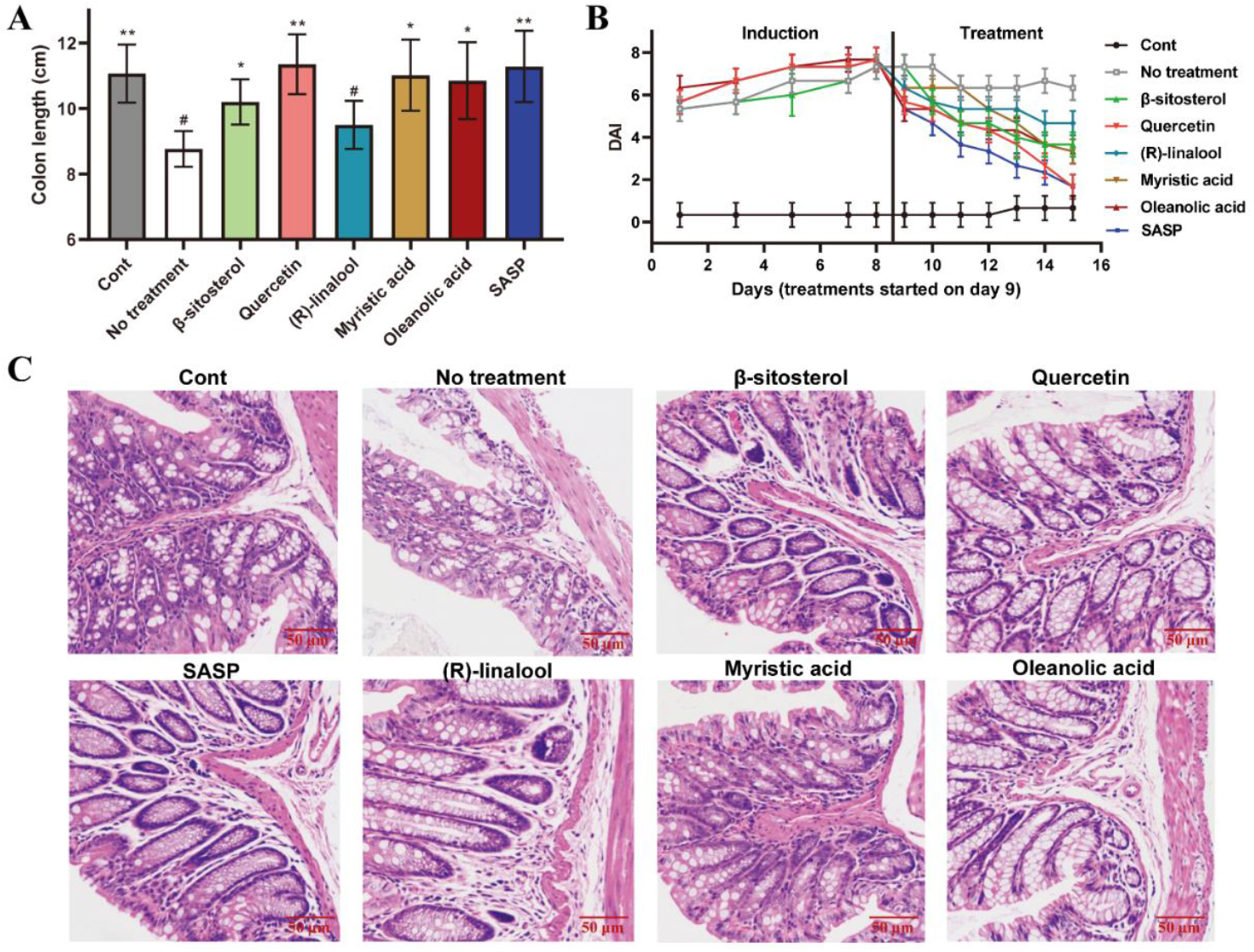
Alleviation of DSS-induced colitis in mice by the 5 bioactive constituents. The 5 constituents, including β-sitosterol (100 mg/kg), quercetin (50 mg/kg), (R)-linalool (10 mg/kg), myristic acid (100 mg/kg) and oleanolic acid (10 mg/kg), were administrated daily to the mice with colitis via gavage for 7 d. SASP (100 mg/kg) was utilized as the positive control. DAI was calculated as the combined score of body weight and stool consistency. A, colon length of the treated mice. B, DAI indicating the severity of colitis. C, H&E staining of the colon tissues. Results are mean ± SD. *Significantly different from the No treatment group (p < 0.05). **Significantly different from the No treatment group (p < 0.001). #Significantly different from the Cont group (p < 0.05).

**Figure. 8.**
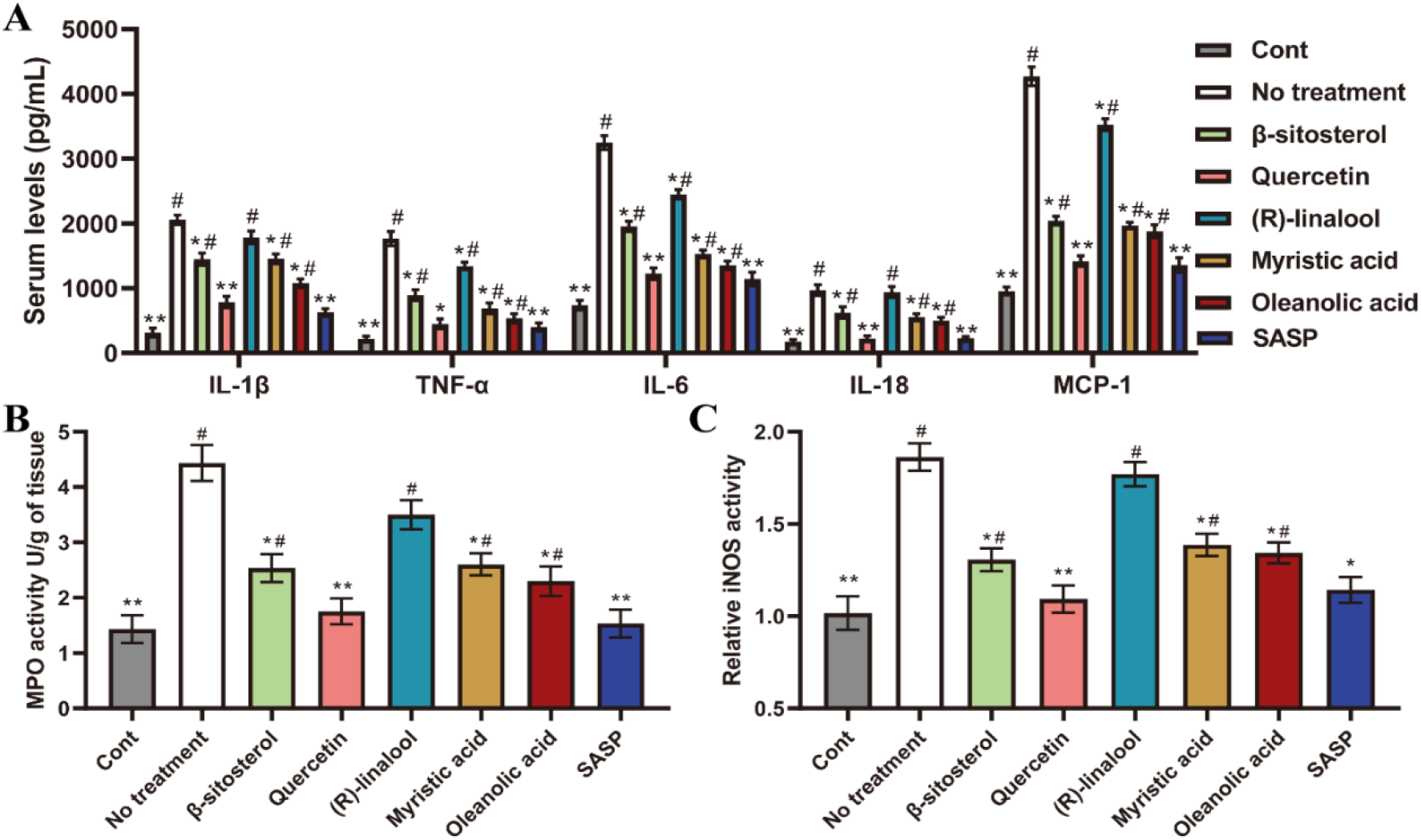
Alleviation of colitis-associated inflammatory responses in mice by the 5 bioactive constituents. The 5 constituents, including β-sitosterol (100 mg/kg), quercetin (50 mg/kg), (R)-linalool (10 mg/kg), myristic acid (100 mg/kg) and oleanolic acid (10 mg/kg), were administrated daily to the mice with colitis via gavage for 7 d. SASP (100 mg/kg) was utilized as the positive control. Inflammatory cytokines in serum were measured by ELISA assays. MPO activity was measured in homogenized colon tissue samples and iNOS activity was evaluated in isolated colon macrophages. A, serum levels of inflammatory cytokines. B, MPO activity in colon tissues. C, iNOS activity in colon macrophages. Results are mean ± SD. *Significantly different from th e No treatment group (p < 0.05). **Significantly different from the No treatment group (p < 0.001). #Significantly different from the Cont group (p < 0.05).

## IV. DISCUSSION

TCM formulas usually consist several herbs, and each herb may have hundreds and thousands of ingredients with bioactivity. In the present study, a DNN was constructed to deal with the complicated situation and utilized for TCM formula recommendations. We presented a model that combined Node2Vec and MLP, which achieved satisfying results on the TCM dataset. The detailed information contained in herbs was constructed into a homogeneous undirected graph, and the Node2vec was adopted for its superior performance in homogeneous graph embedding tasks. The lack of structural prior provided the present construction a general representational ability that could be utilized as a baseline for future network designs. On the other hand, despite the fact that Node2Vec is effective for even large-scale graph embedding task, its approach of exploring the local neighborhood of nodes imposes restrictions on representing the global structure of graph. Therefore, other models can be employed to model the herb-constituent-structure relations with high-complexity in our future work. Currently, the graph embedding models based on auto-encoder (AE), such as VGAE [17], ANE [18] and GALA [19], are effective in modeling the nonlinear structure of graph. As a deep model that specializes in processing graph-type data, the graph neural network (GNN) is able to generate embeddings that retain neighborhood information with any depth. For instance, GAT employed the attention mechanism to strengthen the capability of embedding representation [20], and SDGNN combined traditional GNN with balance and status theories which could effectively generate directed graph embeddings [21]. As a typical work of merging GNN with transformer, Graphormer was known for its considerable performance in graph embedding tasks of microscopic information such as the molecular structure [22]. The present research focused on colitis-related content, as the scale of the data was limited. In order to utilize those sophisticated models to obtain more implicit information related to TCM formulas, the data size of prescriptions should be tremendously expanded so that the herb-herb and target-target interactions can be supplemented to the graph to enhance the efficacy of prescription productions.

Colitis is known as the acute or chronic inflammatory conditions in colon, which can be induced by infections, drugs or various undetermined causes [23], [24]. Many classical TCM formulas, such as Shenling Baizhu powder [25], Fuzi Lizhong pill [26] and Sijunzi decoction [27], have been utilized in clinical practice for the treatment of colitis.

Identification of the major bioactive constituents is critical for the exploration of molecular targets and therapeutic mechanisms of these formulas. In the present study, 5 major constituents were identified among 378 prescriptions through neural network and network pharmacology, and their treatment effects were also verified in mice with DSS-induced colitis. Among the 5 identified constituents, the activities of β-sitosterol [28], quercetin [29], (R)-linalool [30] and oleanolic acid [31] against colitis have already been revealed in previous studies, which also support our observations. However, all these studies revealed the anti-inflammatory and antioxidant potential of these compounds without clarifying the definite protein targets. With the help of the TCM databases and network pharmacology, we were able to uncover the underlying targets and cell signaling pathways, such as the hypoxia-related HIF-1 or inflammatory cytokine IL-17, which will benefit future pharmacological studies on therapeutic natural products for colitis treatment.

Besides the identified bioactive constituents, there are thousands of other ingredients in a TCM formula which target various molecules in different cell signaling pathways. In addition, cell signaling pathways are known to be interact with each other to affect similar biological activities, constructing a complicated interactome network [32]. In this case, the therapeutic outcome of a certain TCM formula could be the additive or synergistic effects of activations or inhibitions of multiple proteins. In the present study, these interactions were overlooked and it was only focused on the anti-inflammatory potential of certain natural compounds identified via the neural network and network pharmacology. The DNN could be updated in future studies by adding more layers to construct the cell-cell and protein-protein interactions in futures studies for generations of more accurate and applicable TCM prescriptions.

In summary, a DNN was constructed based on herb-ingredient-target relations through the augmentation of 378 distinct TCM formulas collected from 204 TCM books or databases for the treatment of colitis to generate high-quality TCM prescriptions. Network pharmacology study of the TCM formulas also revealed the anti-inflammatory potentials of the most frequent ingredients in the formulas. The therapeutic and anti-inflammatory effects of the 5 major bioactive constituents in the intersection between the compounds in top-scored prescription and the most frequent molecules in all prescriptions were verified in mice with DSS-induced colitis, which not only confirmed the rationality of the generated TCM prescription and network pharmacology outputs, but also indicated that these compounds could be utilized in the development of therapeutic agents for colitis.

## Supplementary Materials

Supplemetary materials 1: Augmentated TCM prescriptions. Supplemetary materials 2: Figure S1: Biological pathway (BP), cell component (CC) and Molecular Function (MF) enrichment with the 102 intersected genes between the 10 most frequent molecules and colitis; Table S1: The scoring criteria of colitis Disease Activity Index (DAI); Figure S2: The induction of colitis in mice using dextran sulfate sodium salt (DSS) and treatments with different agents.

## Ehical Approval

The animal experiments were conducted following the protocols approved by the institutional Laboratory Animal Use and Care Committee (IACUC, prot. No. ECUST-2022-0089) and all applicable international and national guidelines for the use and care of animals were strictly followed.

## Data Availability Statement

Data is contained within the article or supplementary materials. The experimental settings, codes and DNN constructions can be found on our Github webpage (https://github.com/AI-HPC-Research-Team/TCM_prescription_generation).

## Acknowledgment

The authors appreciate Yue Zhou from Peng Cheng Laboratory for the technical advice and Zhaoqian Ding, Ying Xie, Tingqian Wang, Linyi Li from East China University of Science and Technology for the collection of TCM prescriptions from books. The research was supported by the Peng Cheng Laboratory and Peng Cheng Cloud-Brain.

